# TSAR, “Thermal Shift Analysis in R”, identifies endogenous molecules that interact with HIV-1 capsid hexamers

**DOI:** 10.1101/2023.11.29.569293

**Authors:** William M. McFadden, Xinlin Gao, Zhijiang Ye, Xin Wen, Zach C. Lorson, Huanchun Zheng, Jessica Fahim, Andres E. Castaner, Karen A. Kirby, Stefan G. Sarafianos

**Affiliations:** Center for ViroScience and Cure, Laboratory of Biochemical Pharmacology, Department of Pediatrics, Emory University School of Medicine, Atlanta, GA; Children’s Healthcare of Atlanta, Atlanta, GA

**Author notes:** These authors contributed equally (William M. McFadden, Xinlin Gao, Zhijiang Ye).

**Keywords:** thermal shift assay, differential scanning fluorimetry, human immunodeficiency virus, capsid, endogenous metabolites, gallic acid, folic acid

## Abstract

The thermal shift assay (TSA) is a versatile biophysical technique for studying protein-ligand interactions *in vitro*. Here, we report a free, open-source software tool, TSAR (“Thermal Shift Analysis in R”), to expedite the analysis of TSA data. The TSAR package incorporates multiple workflows that facilitate TSA analyses, returns publication-ready graphics, and includes an optional graphic user interface. The package is available at https://bioconductor.org/packages/TSAR/. Applying TSAR, we screened two chemical libraries and found multiple molecules that potentially interact *in vitro* with the capsid protein (CA) of human immunodeficiency virus type 1 (HIV-1). First, a library of vitamins exemplifies the different graphic outputs of TSAR, and we report a change in the 50% melting temperature (ΔT_m_) for folic acid-treated CA hexamers (CA_HEX_). Since HIV-1 CA_HEX_ interacts with host-derived acids like inositol hexaphosphate (IP6) or dNTPs, a second library was screened containing 96 organic, acidic metabolites; multiple anionic ligands caused a ΔT_m_ for CA_HEX_. Subsequent investigation of these interactions includes biolayer interferometry, antiviral activity against pseudotyped HIV-1, and endogenous reverse transcriptase assays that were used to validate and investigate the biological impact of these native ligands that thermally-stabilize CA_HEX_. One compound hit, gallic acid, exhibited anti–HIV-1 activity as previously reported, and we show interacts with CA_HEX_ as a potentially novel mechanism. Overall, the TSAR package facilitated quick analysis of TSA data from multiple libraries to help identify a biologically relevant hit, gallic acid, as a molecule that can inhibit HIV-1 replication and targets CA_HEX_.

**Importance:** The TSAR package is freely available (AGPL-3) and is designed for both experienced or new R users, having command-line code for handling large and challenging datasets while also including an optional GUI that enables easy use by non-programmers. This is the first TSA analysis program written in R, a free and open-source language; TSAR simplifies TSA analysis while maintaining diverse visualization options for small-to-large libraries and multidimensional analysis. Additionally, we report multiple endogenous metabolites that potentially interact with the HIV-1 capsid protein hexamers (CA_HEX_) *in vitro*, including folic acid, gallic acid, and multiple others, primarily from the tricarboxylic acid (TCA) cycle. Various methods validate gallic acid interactions with CA_HEX_, leading to a novel suggested mechanism of action, and in-line with previous reports, this metabolite has potential for natural-based treatments of HIV-1. Further, we advance the understanding of the potential mechanism(s) for HIV-1 inhibition by gallic acid treatment.

## Introduction

The thermal shift assay (TSA), alternatively “differential scanning fluorimetry” (DSF), is an *in vitro* technique to study the thermal stability of a protein or biological complex with multiple applications (1–4). During TSA experiments, samples are heated at a constant rate to track the denaturation of protein(s) of interest (Supplementary Figure 1A). Proteins that contain internal aromatic residues, like tryptophan, can be tracked without a dye by measuring the change in residue fluorescence (4); however, TSA experiments often utilize dyes that fluoresce in hydrophobic environments, like SYPRO™ Orange, to follow the exposure of the internal, hydrophobic residues as denaturation progresses (2, 5). Plotting the temperature compared to the fluorescence reading gives an individual melting profile for the protein or complex measured. The melting curve of uncharacterized proteins can be unpredictable, and testing of various dyes can help validate the experiment (6). For folded and globular proteins, hydrophobic residues are typically internal and inaccessible to the solvent, making TSA a useful tool to assess if a batch of proteins is properly folded and consistently prepared (7, 8).

For most samples that are properly folded, the thermal profile is sigmoidal, and a melting temperature (T_m_) value can be calculated, representing the temperature where 50% of the protein is unfolded (2, 7) (Supplementary Figure 1A). Curves that are not properly sigmoidal will need to be omitted or assessed independently for melting temperature(s). The T_m_ value is used to compare the thermal stability of proteins under various conditions, such as selecting optimal storage buffers and salts or for comparing the stability of mutant proteins (9, 10). Of note, intrinsically disordered proteins do not typically have a sigmoidal curve and lack T_m_ value at room temperature, as they are unfolded by definition (11, 12). TSA is also useful for screening chemical libraries to identify potential binding partners, since protein•ligand and protein•protein interactions that form additional intramolecular bonds and thus require more energy, or a higher temperature, to denature the sample compared to an unliganded sample (1, 2, 13–15). The “thermal shift”, determined by the change in the T_m_ (ΔT_m_), is determined by calculating the difference between the T_m_ of treated and untreated sample(s). Thus, TSA is used as a proxy for *in vitro* ligand binding to purified protein targets within screens for molecular interactors such as inhibitors, natural ligands, and cofactors.

The TSA technique is used extensively in labs that focus on biological research, as it requires only a qPCR machine for data acquisition and one of many commercially-available dyes, like SYPRO™ Orange, once a protein or biological complex is purified (2, 6). The required volume and concentration of the sample are quite low, typically under 25 μL and below 20 μM per sample, although multiple replicates are needed for comparison and statistical analysis (1–4). Some popular TSA analysis tools are Protein Thermal Shift™ Software, Serial Explorer in MATLAB, and open-source workflows provided through the Konstanz Information Miner (16), as well as some other free tools (17, 18). While both Protein Thermal Shift™ and MATLAB provide high-throughput analysis, they are proprietary and require additional costs. Despite other tools being free and open-source, they often offer limited visualization or analysis options. Given the need to visualize the data in multiple dimensions and enhance accessibility of analysis tools, we developed a software package written in R (19), a free programming language extensively used for biological data analysis. ‘TSAR’, short for Thermal Shift Analysis in R, is a free and open-source R package (Figure 1). TSAR is hosted by *Bioconductor* and distributed under an AGPL-3 license that offers a workflow as both command-line functions and an interactive ShinyR graphic user interface (GUI) dashboard (20, 21). TSAR melting curve analysis allows users to select Boltzmann-fit or derivative-modeling of data to identify either T_m_B (Boltzmann-fit) or T_m_D (derivative-model) from melting curves that are smoothened with beta-knot fitting (Supplementary Figure S1B-C) (22, 23). The TSAR package is a complete suite of tools from initial data processing, T_m_ identification, multidimensional comparisons, and visualization of various publication-quality graphics from individual experiments to large screens with any number of replicates (Figure 1B-E). TSAR is a well-documented package and has many custom settings for users experienced in R, while it also incorporates wrapping functions and a ShinyR dashboard to minimize coding for easy use and quick implementation (20, 24). Overall, TSAR aims to simplify TSA analysis yet produce diverse visualizations.

**Figure 1:**
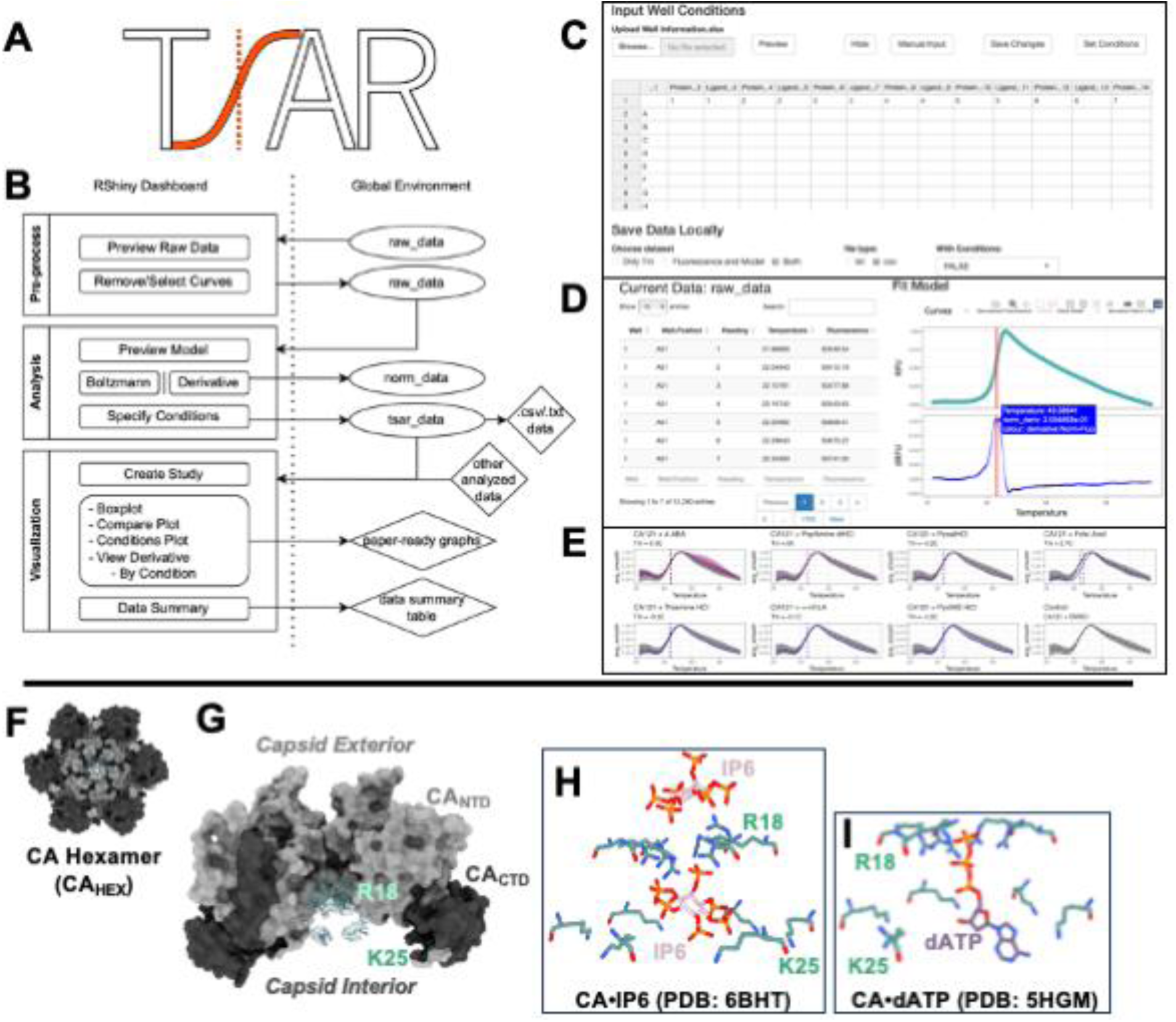
TSAR (*Thermal Shift Analysis in R*) package overview and example protein target. **A)** TSAR Logo. **B)** Workflow for TSAR, supporting data import and export at various stages. **C-E)** Screenshots of the TSAR program. **C)** shows the Shiny GUI for importing data, **D)** shows the interactive data analysis that TSAR performs, and **E)** is an example of a partial output from a library screen. **F)** Structure of WT HIV-1 CA_HEX_, visualized with ChimeraX (PDB: 4XFX)(26), **G)** Partial side view of (F), showing the cationic central pore with Arg18 and Lys25 in stick representation. **H)** Structure of IP6 in the CA_HEX_ central pore (PDB: 6BHT)(48). **I)** Structure of dATP in the CA_HEX_ central pore (PDB: 5HGM)(47).

Here, we perform two example screens using TSAR to identify potential binding partners to the human immunodeficiency virus type 1 (HIV-1) capsid protein (CA). The HIV-1 capsid core, made of over 1500 copies of CA, is an essential macromolecular structure within the mature virus that serves many roles during infection (25–28). The capsid core is the reaction chamber for reverse transcription, it is a shield to hide the viral genome from cellular immune sensors, and it is the vehicle that traffics the HIV-1 genetic material through the cytoplasm to, and likely through, the nuclear pore complex (28–37). As it is essential in numerous steps of HIV-1 replication, investigating the biology of the capsid is pivotal in the design of next-generation antiretroviral therapies (ART) (38–40). In fact, when the CA-targeting compound Lenacapavir (LEN, formerly GS-6207) was approved for use in highly-treatment experienced patients in 2022, CA became the first new target for a clinically-used ART class in over a decade (38, 41). Recently, LEN was additionally approved for use as pre-exposure prophylaxis (PrEP), reviewed in (42). Thus, understanding the biology of this clinically-relevant target is of high importance.

One reason that the capsid core is a promising antiretroviral target is that CA is a highly conserved domain and genetically fragile protein; the kinetics of CA•CA interactions are finely tuned by evolution for balancing cellular trafficking and genome release, such that both increasing and decreasing capsid stability has a negative impact on viral fitness (9, 28, 40, 43–46). The mature HIV-1 core is comprised of primarily hexameric (CA_HEX_) multimers of CA (Figure 1F), which form a central pore with multiple basic residues like Arg18 and Lys25 (Figure 1G). In fact, reports have demonstrated that CA_HEX_ interacts with host-derived acidic metabolites like dNTPs and inositol hexaphosphate (IP6) at this pore to facilitate efficient reverse transcription and promote stable capsid assemblies (Figure 1H-I) (28, 32, 47–51). Considering that the mature capsid navigates through the cytoplasm during infection (27, 42, 47), in addition to reports that HIV-1 infection modulates cellular metabolism both in the presence and absence of ART (52–54), we hypothesized that there might be additional endogenous interactions with other common protein cofactors or naturally-present acidic small molecules. Thus, two example screens are presented to identify potential interactions of vitamins or anionic metabolites with purified CA_HEX_ (55). First, a small library of essential vitamins that are enzymatic cofactors finds a potential *in vitro* interaction with folic acid (vitamin B9). Second, a library of 96 acidic metabolites finds many anionic compounds that increase CA_HEX_ T_m_. This library contained internal control of IP6 (“phytic acid”) that had the largest ΔT_m_; thermally-stabilizing hits include gallic acid, xanthurenic acid, metabolites from the tricarboxylic acid (TCA) cycle, and others. Both *in vitro* activity and cell-based antiviral activity were tested for multiple hits, and gallic acid routinely affected HIV-1 replication or *in vitro* enzymatic activity when the mature capsid was present. Gallic acid has been shown to have antiviral activity against HIV-1 previously, however these studies point to targets like the reverse transcriptase (RT) enzyme (56–58); our analysis suggests an additional role for capsid core that encapsulates the reverse transcription reaction. These results provide example workflows for the TSAR package, a tool that can be broadly used in protein biology, and the screens presented here have interesting implications for the stability of HIV-1 capsid in the context of cellular metabolism.

## Results

### TSAR Package Accuracy

To assess the accuracy of TSAR, we found the T_m_D and T_m_B return similar values (Supplementary Figure S1C). We benchmarked the determined T_m_ and ΔT_m_ shift from TSAR to those from Protein Thermal Shift Software for the 12-component vitamin library. Using the Protein Thermal Shift Software as the baseline value, TSAR T_m_D error is centered at 0.048 with a variance of 0.063 degree Celsius (Supplementary Figure S1D). Meanwhile, TSAR T_m_B error is centered at 1.029 with a variance of 0.634 degree Celsius (Supplementary Figure S1E). While that error is relatively greater than ΔT_m_D, the error is systematic where computing ΔT_m_B by using DMSO as the baseline for ΔT_m_, errors centering at 0.376 °C (Supplementary Figure S1F-G). Thus, we set T_m_D as the default modeling method in TSAR.

### Vitamin library exemplifies TSAR output types

Testing the vitamin library, with IP6 as a positive control, yielded significant result for folic acid with a ΔT_m_D of 3.04 °C (Figure 2). Figure 2 exemplifies three methods of visualization by TSAR: individual melting curves with an average (Figure 2A), compare averaged melting curves (Figure 2B), and boxplots to show the average ΔT_m_ (Figure 2C). Derivative plots, which show T_m_ as a maximum, are visualized also (Supplementary Figure 2A-C).

**Figure 2:**
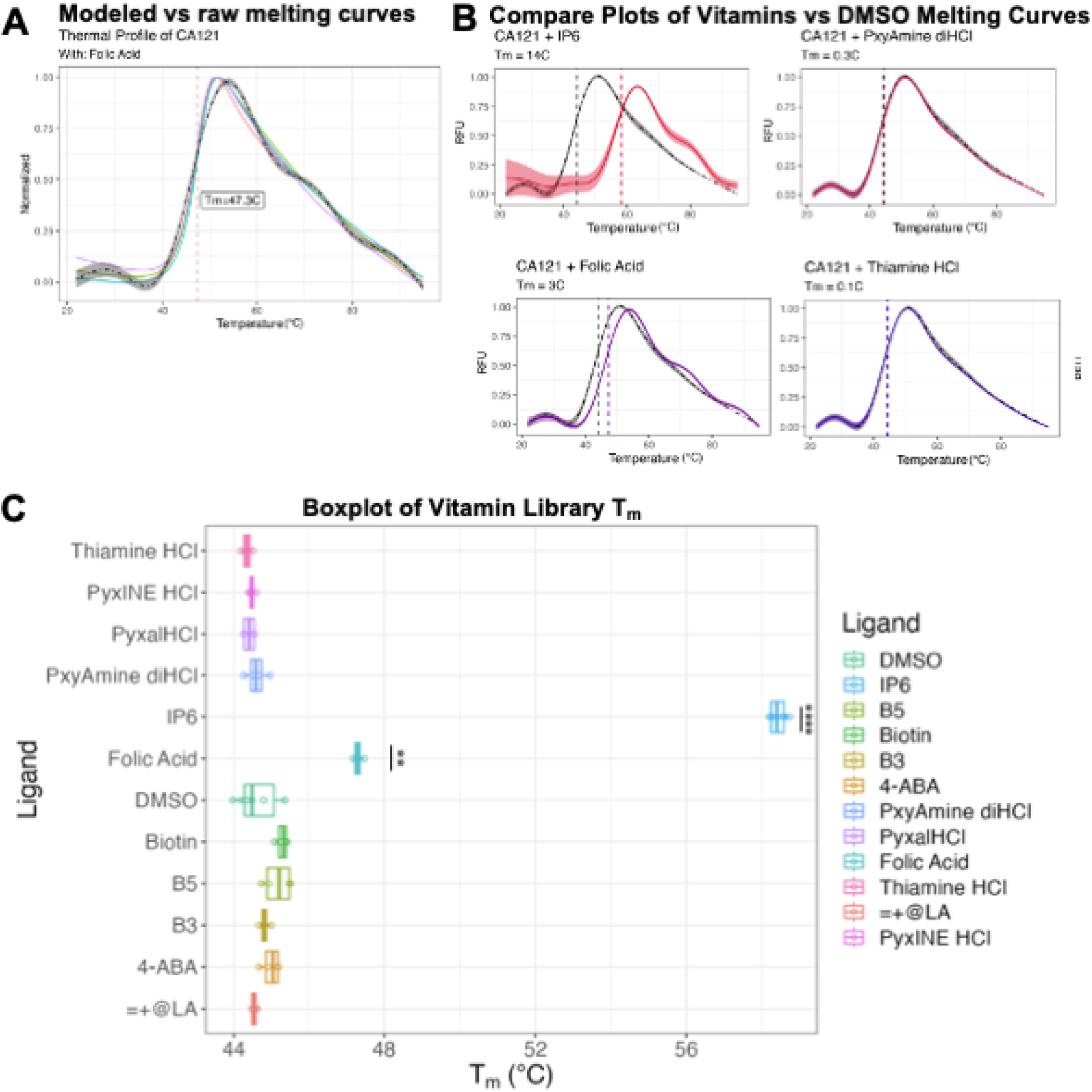
Example TSAR outputs and screening of CA_HEX_ binding to a library of vitamins. **A)** Folic acid thermal melting profiles by individual sample (n = 4). Each well is shown as a different color. Y is normalized relative fluorescence units (RFU). The grey line represents the modeled average between all qPCR wells **B)** Compare plot further showcases ΔT_m_ interactions as averaged RFU over temperature (°C), compared to DMSO control (grey). Only IP6 and folic acid yield significant T_m_ change: IP6 ΔT_m_D=13.75℃ (p<5E-11), folic acid ΔT_m_D=3.04℃ (p<5E-5). C) Screening vitamin library at 50 μM with CA121 (CA_HEX_) at 7.5 μM. Compared to DMSO vehicle.

Testing at 50 μM, 25 μM, and 7.5 μM compound, the first derivative comparison graphs demonstrate that the ΔT_m_ of folic acid with CA121 is dose dependent (Supplementary figure 2A-D). At 7.5 μM, protein unfolding curves are distinguishable from DMSO control, but ΔT_m_ experiences greater variance. Conducting one-tailed, paired T-tests show that all treatment conditions are significantly different from the DMSO control, with 50 μM and 25 μM folic acid (p≤5E-5), and 7.5 μM (p≤5E-3) (Supplementary Figure 2D). Structures of IP6 and Folic acid shown (Supplementary Figure 2E). When breaking the CA_HEX_ into CA monomers using TCEP (Supplementary Figure 2F), we find a loss of ΔT_m_ for both the IP6 control and folic acid, indicating the importance of the multimeric CA (Supplementary figure 2G-H).

### Multiple acidic metabolites cause a ΔT_m_ for CA_HEX_

The second library contained 96 anionic compounds that are part of various metabolic pathways and included an internal control of IP6. As expected, 25 µM IP6 provides an extremely strong ΔT_m_ of 12°C, but multiple other metabolites also had statistically significant ΔT_m_ values (Figure 3A-C, Supplementary Figure 3). Initial hits included gallic acid (ΔT_m_=6.37°C), multiple TCA cycle metabolites like ⍺-ketoglutaric acid (⍺Kg, ΔT_m_=5.79°C), oxaloacetic acid (ΔT_m_=4.99°C), citric acid (ΔT_m_=4.10°C), and others like xanthurenic acid (ΔT_m_=2.28°C) and indole-3-pyruvic acid (ΔT_m_=1.65°C).

**Figure 3:**
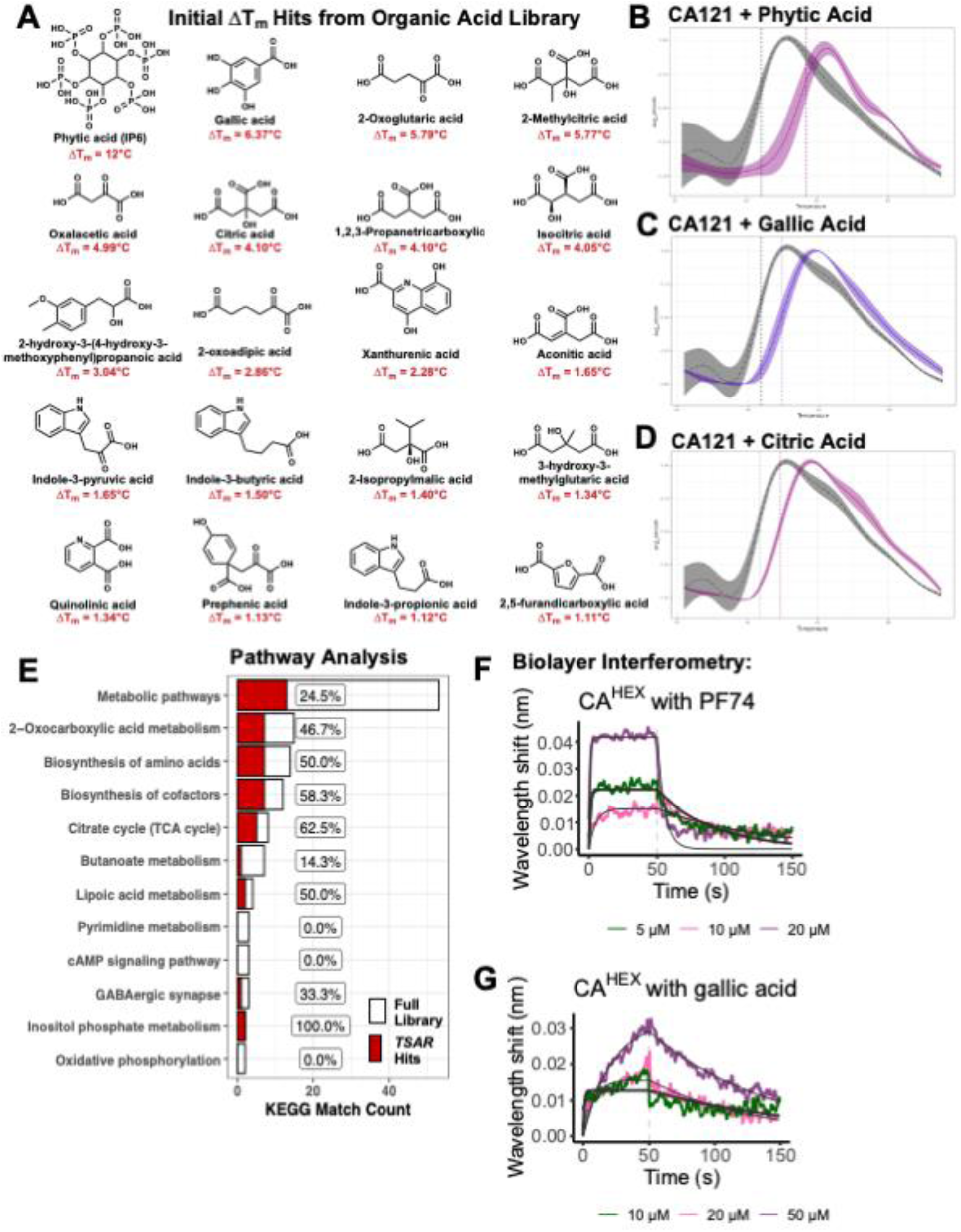
TSA of organic metabolic acids find many potential interactions with CA_HEX_. **A)** Top 20 hits from the 96-compound library in 50 mM Tris (pH 8.0). 25 µM compound and 7.5 µM CA_HEX_ (CA121). Made in ChemDraw (V19.0). See Supplementary Figure 3. **B-C)** TSAR compare plots to show the large ΔT_m_ caused by **B)** phytic acid (IP6, internal control), **C)** gallic acid, and **D)** citric acid. **E)** *H. sapiens* pathway matches to metabolites in the library (in white) or those that cause ΔT_m_ determined by TSAR (in red) in the KEGG Database (59). **F & G)** Biolayer interferometry (BLI) validates interaction of CA_HEX:HIS_ with **F)** PF74 or **G)** gallic acid.

KEGG analysis mapped 53 metabolites in the acidic library to *H. sapiens* pathways (59), and of them, over 60% of the represented TCA cycle metabolites caused a ΔT_m_ (5 hits of 8 possible, 5/8), especially those involved in the synthesis of ⍺Kg (Figure 3E, Supplementary Figure 4A). Other pathways with 50% or more matched metabolites that cause a ΔT_m_ at 25 µM include amino acid biosynthesis (50%, 7/14) especially for tryptophan biosynthesis (66.7%, 4/6), cofactor biosynthesis (58%, 7/12), lipoic acid metabolism (50%, 2/4), and inositol phosphate metabolism (100%, 2/2) (Figure 3E).

To validate the binding of gallic acid to CA_HEX_, we used BLI to determine binding kinetics. We used a HIS-tagged CA_HEX_ (CA_HEX:HIS_) to immobilize the proteins to the HIS-1K biosensor (60). Using blocking buffer that contained Bovine Serum Albumin (BSA), which worked well for other experiments (60–62) including compounds that bind to the phenylalanine-glycine (FG) dipeptide binding site in CA_HEX_ (28, 60), did not return kinetic binding curves that could be analyzed (data not shown). Thus, we optimized the blocking buffer by switching BSA to lysozyme, as BSA is reported to interact with many metabolites including gallic acid (63, 64). We find PF74 binds to CA_HEX:HIS_ with a K_D_ of 4.41±0.16 µM, *k_on_* of 4.45E+03±1.44E+02 M^-1^s^-1^, and *k_off_* of 1.96E-02±3.32E-04 s^-1^ (Figure 3F), slightly different than with the BSA-containing buffer (60, 65), and Gallic acid binds with K_D_ of 22.3±0.64 µM, *k_on_* of 4.02E+02±9.58E+00 M^-1^s^-1^, and *k_off_* of 8.96E-03±1.42E-04 s^-1^ (Figure 3G). This confirms binding of these ligands to CA_HEX_.

### Gallic acid binds CA_HEX_ and has antiviral activity

We first checked the effect of adding these native metabolites in excess would influence HIV-1 infection. No compound was cytotoxic at 100 µM, except for the antiretroviral control PF74 with a 50% cytotoxic concentration (CC_50_) at 75.29 µM (14, 28, 40, 66). PF74 serves as a positive control with a 50% effective concentration (EC_50_) of 0.6±0.1 µM, aligning with literature values (14, 28, 40, 66). Notably, gallic acid had antiviral activity with an EC_50_ of 19.2±0.1 µM for pseudotyped HIV-1_NL4-3ΔEnv_. This inhibition aligns with other reports of gallic acid antiretroviral activity (56–58, 67).

While the PF74 control did increase CA assembly rate as reported, no metabolite tested increased WT CA assembly in an assay that measures rates of assembly using turbidity (Absorbance at 350 nm (A_350_)) as a proxy for lattice assembly (14, 68, 69) (Supplementary Figure 4B).

We further employed the ERT assay that tests the activity of RT within purified, native capsid cores stabilized with IP6 (32, 49). This determines the effect on early, intermediate, and late stages of reverse transcription (reported in (32, 49)), to systematically probe the inhibition of RT. The early steps are somewhat impacted by PF74 treatment, but impacts intermediate and late stages more, as damaging the capsid decreases the reaction efficiency of RT (49). Gallic acid has increasing inhibition as the reaction proceeds: no determined effect on early reverse transcription, a modest effect on intermediate steps (intermediate IC_50_=17.21 µM), and a significant effect on late step products (late IC_50_=3.64 µM). Additionally, the metabolite xanthurenic acid consistently impaired RT from early to late events (IC_50_s = 9.65 µM - 11.99 µM). Other metabolites tested for ERT and antiviral activity had no effect, including folic acid, oxalacetic acid, citric acid, isocitric acid, tricarballylic acid, 2-methylcitric acid, ⍺Kg, and (Z)-aconitic acid (data not shown).

To further probe if the polymerase activity of RT is inhibited by gallic acid, we used a reported primer-extension assay applying purified enzyme to a short template and treated with inhibitors (70, 71). While RT inhibitors doravirine (DOR) and islatravir (EFdA) impair primer extension (71), up to 1000 µM of gallic acid had no effect on the polymerase activity of RT, similar to the other CA inhibitors tested, PF74 and LEN, that do not target RT (Supplementary Figure 4C).

## Discussion

Here, we performed TSA screens to identify potential protein•ligand interactions that was conducted with TSAR and found that multiple metabolites stabilize HIV-1 CA_HEX_ *in vitro*. TSAR is a novel package written in the R language, which is a tool used by many biological research labs and in higher-education coursework (72). There are other software tools for TSA data analysis, but they are costly or have limited visualization options. Meltdown is a similar tool and is a free and user-friendly python package (18), however, TSAR uniquely provides T_m_ estimation using the boltzman-fit method and includes a GUI dashboard. MoltenProt is a free and useful online tool but requires additional visualization software for high-resolution graphics (17); TSAR is free for academics and can output publication-ready graphics.

There are reports identifying endogenous metabolites that interact with mature CA_HEX_ being critical cofactors in HIV-1 replication, such as IP6 that is important for capsid stability as well as dNTPs that are needed for genome replication (28–31, 47, 48). TSAs alone do not establish the virological significance of these metabolites, only that stabilizing interactions are observed *in vitro* for CA_HEX_. By investigating the TSA hits, our studies that included gallic acid find antiviral activity and inhibition of RT activity, aligned with some prior reports about the inhibition of reverse transcription activity and HIV-1 infection by gallic acid (56–58, 67). Using ERT assays where is RT inside the capsid core (32, 49), the intermediate and late reverse transcription products were inhibited by gallic acid, but not if RT is alone in the *in vitro* primer-extension assay that tests its polymerase activity (70, 71). CA_HEX_ are involved dNTP import that fuel RT; it is shown that blocking the capsid pore with the flat, inorganic acid hexacarboxybenzene can impact reverse transcription (47, 49, 73). Attempts to solve the atomic structure of CA with gallic acid by X-ray crystallography were not successful (data not shown). Other functions of RT may be impacted, such as RNase H activity or metal ion coordination (67, 74), as gallic acid is a divalent ion chelator (75). If the enzymatic target is RT, as proposed by others (56, 57), our results indicate there is a relationship with gallic acid, a non-nucleoside molecule that, in theory, would need to traverse the mature capsid core and reach RT or impair its function distant from the enzyme.

We did not anticipate that supplementing cells with endogenous metabolites would impair viral fitness, as most metabolites are readily converted to other molecules by cellular enzymes, and the virus replicates within biologically active cells that typically contain these metabolites. No observable effect was the case for almost all TSA hits, except for gallic acid. Gallic acid is a secondary metabolite typically acquired through diet or gut microbiota; concentrations in human cells are generally low and there are reports of gallic acid cytotoxicity in some cell-types (75, 76). Gallic acid has been shown to have antiviral activity for multiple viruses, including HIV-1, influenza, and encephalomyocarditis virus (56, 57, 76–78), indicating a complex and likely multifaceted mechanism. Gallic acid is also a sticky molecule, and is known to interact with many cellular and viral proteins, including a report that used surface plasmon resonance and found interactions with HIV-1 proteins CA, RT and Gag (57, 63, 64, 79–81). ERT assays do not find an impact on *in vitro* RT products by treatment with most of the tested metabolites, except gallic acid and xanthurenic acid. Xanthurenic acid is a metabolite involved in the tryptophan oxidation pathway with concentrations reportedly decreased in ART-naïve participants living with HIV-1 (54, 82); we found it impacts *in vitro* ERT but not an observable antiretroviral effect *in vivo* (Figure 4A-D).

**Figure 4:**
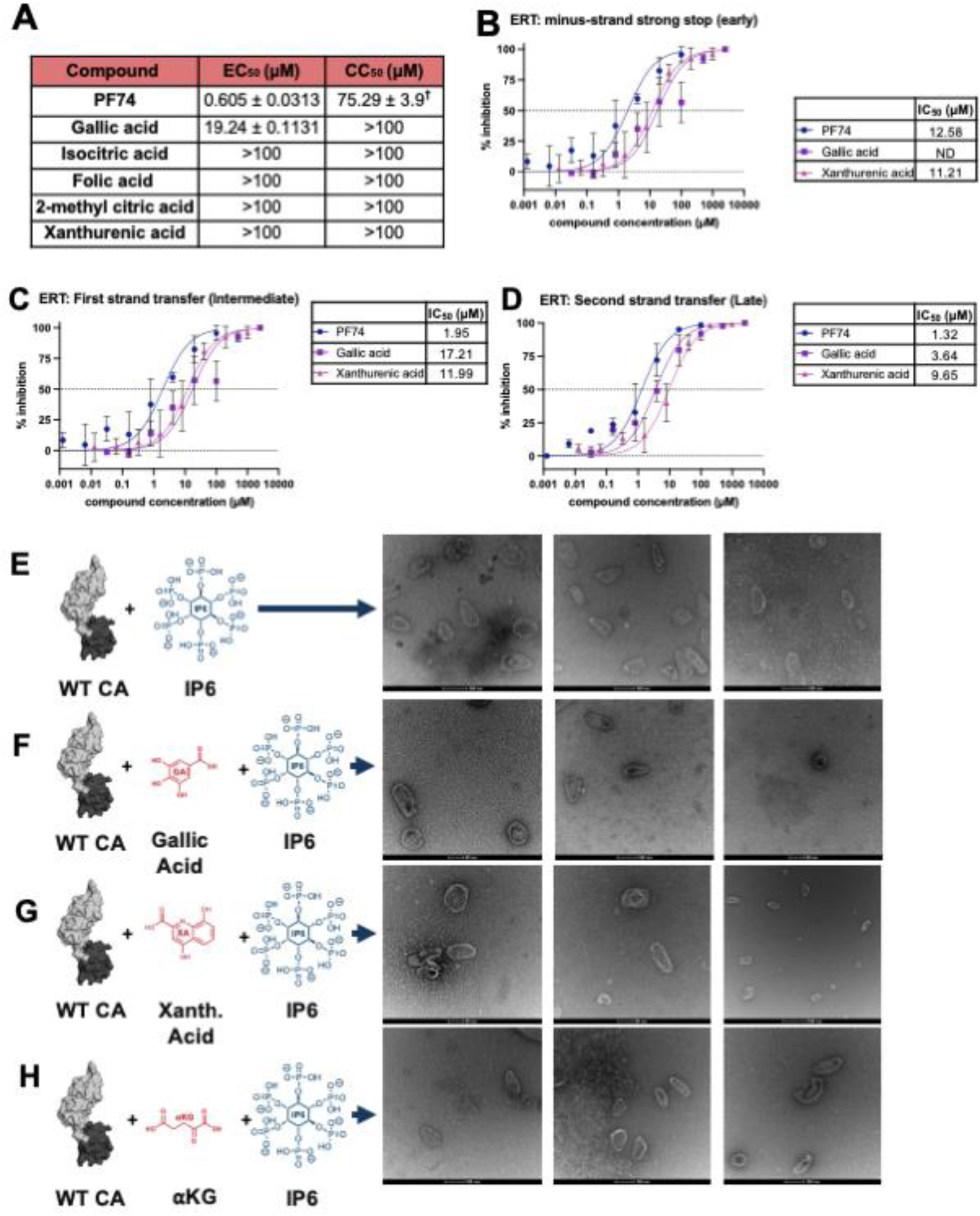
Testing the virological effects of gallic acid and other CA_HEX_-stabilizing metabolites on HIV-1 replication. **A)** Antiviral activity (EC_50_) using HIV-1_NL4-3_ pseudotyped virus in TZM-GFP cells and cytotoxicity (CC_50_) of select compounds. ^†^PF74 CC_50_ from (14). **B-D)** Endogenous reverse transcription (ERT), as previously described (32, 49), takes place inside an IP6-stabilized capsids requiring both CA and RT function: **B)** early, **C)** intermediate, and **D)** late RT products are determined by qPCR after incubation with compounds to determine inhibitory concentrations (IC_50_), compared to DMSO control. ND=Not Determined. **E-H)** Negative stain transmission electron microscopy (TEM) of *in vitro* assembled capsid-like particles (CLPs). Monomeric CA mixed with an excess of IP6 form conical assemblies, from (48, 50). **E)** IP6 alone, or IP6 plus **F)** gallic acid, **G)** xanthurenic Acid, or **H)** or 2-oxoglutaric acid (⍺KG) are added to WT CA. Scale varies. Chemical structures made in ChemDraw (V.19.0) and protein structures in Chimera X (PDB: 4XFX)(26).

Numerous studies have investigated the requisites for and changes in metabolism during HIV-1 infection (53, 54, 82, 83). Of note, a report found that cells lacking mitochondria inhibit HIV-1 post-entry and pre-integration (83), a phenotype typical with dysfunctional capsid; another study found increased HIV-1 infection when the TCA cycle was highly active (53). While only the TSAs showed an impact on CA thermal stability and not an effect in other virological assays, the assays were tested with excess metabolites and not in the absence, which may have a different effect. The role and relevance of these endogenous molecules, both the vitamin folic acid and the organic acidic metabolites, in HIV-1 biology remains a topic of investigation, but the findings of our screen reported here indicate the ability for CA_HEX_ to form many stabilizing interactions with endogenous polyanions. We report here that some CA-stabilizing molecules, identified by TSAR, can impair HIV-1 reverse transcription inside capsid cores, and we further the mechanistic knowledge of gallic acid antiretroviral activity.

## Methods

### Distribution and Package Information

TSAR is an open-source package written in R (Version (V)≥4.3) (19), distributed under an AGPL-3 license in the peer-reviewed *Bioconductor* R package repository (V≥3.18) (21, 84) at https://bioconductor.org/packages/TSAR. TSAR was coded using RStudio (85) and devtools (86). TSAR contains well-documented functions and multiple vignettes (long-form documentation) (24). TSAR depends on tidyverse packages (87) as well as: jsonlite (88), openxlsx (89), mgcv (90–92), and minpack.lm (93) for analysis, and shiny (20), shinyWidgets (94), plotly (95), shinyjs (96), rhandsontable (97), and ggpubr (98) packages for the GUI and data visualizations.

### Package Structure

TSAR has three individual operating dashboards, “Data Preprocessing”, “Data Analysis”, and “Data Visualization”, each with one command line call (Figure 1B). The GUI dashboard is optional, as TSAR also allows command-line functions. The package also enables “Data Visualization”-only, if suppled with pre-determined T_m_ values to use TSAR to visualize previously analyzed data. Mathematical modeling in TSAR relies on both generalized additive models (GAM) cubic spline with beta-knots or Boltzmann fitting to compute T_m_D and T_m_B values (22, 23, 90, 92, 99, 100). See package documentation in TSAR vignettes (24).

### Statistics and Analysis

TSAR utilizes GAM from the mgcv package to capture the derivatives of the curve and the inflection point, T_m_D (90–92). Alternatively, TSAR utilizes non-linear modeling from the minpack.lm package (93) in R to capture Boltzmann curves with machine-learned minimum and maximum, T_m_B.

T_m_B uses a kinetic equation describing the statistical behavior of a system of particles (99); it is applied widely towards sigmoidal behaviors. Using parameter-based mathematical modeling, individual melting curves are fit through a 5-parameter function (101) (Supplementary Figure S1B).

T_m_D, representing T_m_ values estimated using a derivative-model, utilizes smoothing to remove noises from data and produces a polynomial function closely representing the trend and shape of raw data. T_m_s are then derived through the calculation of first derivatives and location inflection point (101). Both methods are used in extant research protocols where T_m_B are less affected by trivial fluctuation and T_m_D captures all levels of variations throughout the denaturation process.

### Protein Purification

The CA_HEX_ with A14C/E45C/W184A/M185A mutations (CA121) were cloned into a pET11a expression plasmid (55), kindly provided by Dr. Owen Pornillos (University of Utah). The expression and purification of CA121 were carried out in *E. coli* BL21(DE3)RIL, and purification and cross-linking was performed as previously described to generate individual, soluble, and disulfide-stabilized CA_HEX_ (14, 55). A C-terminal His-tag was cloned onto the CA121 construct (Genscript) for Biolayer Interferometry (BLI) experiments (60, 61), and this construct was expressed *E. coli* NiCo21(DE3) and lysed and cross-linked as in (55), but purified by Ni^2+^ chromatography before crosslinking and a final size-exclusion chromatography (SEC) step to separate cross-linked CA_HEX_ from monomeric CA.

Additionally, monomeric full-length, wild-type CA (WT CA) was cloned into a pET24a expression plasmid, kindly provided by Dr. Robert Dick (Emory University). The expression of WT CA was performed in *E. coli* NiCo21(DE3) and purification involved ammonium sulfate precipitation followed by desalting, cation-anion subtractive chromatography, and SEC as previously described (50).

HIV-1 RT was expressed and purified as previously described (70). Briefly, pRT6H-PROT (102) was expressed in *E. coli* BL21(DE3)RIL and purified by Ni^2+^ chromatography.

### Compounds

The vitamin library of chemicals were purchased as the Vitamins Kit (Sigma Aldrich, V1-1KT), resuspended in DMSO to a final concentration of 10-50 mM and stored at −20°C. Compounds included: D-pantothenic acid hemicalcium salt (B5), biotin, nicotinamide (B3), 4-aminobenzoic acid (4-ABA), pyridoxamine dihydrochloride (PyxAmine diHCl), pyridoxine hydrochloride (PyxINE HCl), thiamine hydrochloride (Thiamine HCl), folic acid (B9), (±)-α-lipoic acid (a-LA), pyridoxal hydrochloride (Pyxal HCl). 50% phytic acid solution (TCI) was used for IP6 positive controls. The Organic Acid Metabolite Library of Standards (OAMLS) was used (IROA Technologies), and DMSO was added to a final concentration of 5 mM. Phytic Acid is within OAMLS and served as an internal control. After initial screening, select chemicals were purchased: gallic acid [HY-N0523], xanthurenic acid [HY-W014666], dihydroferulic acid [HY-N7080] (similar to 2-hydroxy-3-(4-hydroxy-3-methoxyphenyl)propanoic acid), indole-3-pyruvic acid [HY-W028393], oxalacetic acid [HY-W010382], DL-isocitric acid trisodium salt [HY-W009362], citric acid [HY-N1428], tricarballylic acid [HY-W020215], 2-methylcitric acid [HY-113371], ⍺-ketoglutaric acid [HY-W013636], and (Z)-aconitic acid [HY-W016814] (MedChemExpress). PF74 powder [SML0835] was purchased from Sigma-Aldrich; EFdA-TP tetraammonium [HY-138561A] and Doravirine [HY-16767] were purchased from MedChemExpress. LEN (HRP-20266) was kindly provided by the NIH HIV Reagents Program. dNTPs were purchased from Promega. Oligonucleotides were chemically synthesized and purchased from Integrated DNA Technologies^TM^.

### Thermal Shift Assay (TSA)

TSAs were conducted using QuantStudio 3 qPCR machine (Thermo Fisher Scientific); the final reaction was 20 µL per well of 7.5 µM CA in 50 mM Tris (pH 8.0) and 1x SYPRO™ Orange Protein Gel Stain (Life Technologies) after incubation with compounds for 30 min on ice. Fluorescence was measured while heating consistently increased 0.2°C/10s from 25–95°C (6, 9, 14, 103). For initial screening, CA121 was incubated with 25 µM of each compound. Some previous reports use 50 mM sodium phosphate buffer (pH 8.0), however, the polyanionic phosphates can potentially compete acidic metabolites for binding to the CA_HEX_, thus, we opted for 50 mM Tris (pH 8.0) instead. Thermal profiles were assessed using TSAR or Protein Thermal Shift Software (V1.4, Applied Biosystems). The compounds that induced a significant ΔT_m_ (>1°C) at 25 µM underwent further testing in a dose-response manner under concentrations of 7.5 µM and 50 µM.

Lastly, vitamins leading to significant thermal shifts were also tested in either WT CA monomers or in the presence of the reducing agent tris(2-carboxyethyl)phosphine (TCEP). TCEP breaks the CA121 disulfide-stabilizing bonds, likely affecting the stability of pockets formed by multiple CA monomers. Comparing the fluorescence profile of TSA samples with and without TCEP provides insight into the potential interactions of a compound with monomeric versus hexameric CA.

Metabolic pathway analysis was conducted on the KEGG webserver (59). The KEGG/CSI IDs were provided by the OMAL library manufacturer, and those without matches were manually searched. Of the 96 compounds, 53 were successfully matched to *H. sapiens* pathways, including the 13 of the 20 ΔT_m_ hits. Hits not mapped to human pathways are: gallic acid, 1,2,3,-propanetricarboxylic acid, vanillactic acid, 3-Hydroxy-3-methylglutaric acid, indole-3-butyric acid, indole-3-propionic acid, and 2,5-furandicarboxylic acid.

### Biolayer Interferometry (BLI)

The BLI protocol was adapted from (60, 61) with minor modifications. Briefly, CA_HEX:HIS_ was immobilized onto a HIS1K biosensor at 100 µg/mL for 600 seconds. Loaded probes were equilibrated for 120 seconds before being exposed to a series of association and dissociation phases. Starting at the lowest analyte concentration in the series, CA_HEX:HIS_ was dipped into a well containing analyte (gallic acid, PF74 (60, 65), or vehicle control) for 100 seconds at concentrations of 5, 10, 20, or 50 µM. Probes were then allowed to dissociate in a well containing blocking buffer for 100 seconds. Blocking buffer contained: 50 mM Tris (pH 8.0), 1 mg/mL lysozyme, 0.6 M Sucrose, and 20 mM imidazole. Three experiments were initially analyzed using the Octet software where replicates were combined, and double reference subtraction was applied. Furthermore, the Y-axis was aligned to the baseline step, the inter-step correction was set to baseline at time 0 s, and Savitzky-Golay Filtering was applied.

### Antiviral Activity and Cytotoxicity

Antiviral and cytotoxicity was determined as previously reported (14, 104). Pseudotyped HIV-1 virions (VSV-G-NL4-3ΔEnv) (105) were produced in HEK-293/17 cells that were cultured in DMEM (Corning), and supplemented with 2 mM L-glutamine, 10% FBS, and 100 U/mL penicillin/streptomycin and harvested 48 hpi. Infection was determined in TZM-GFP cells, a reporter-cell line that expresses GFP induced by Tat and regulated by Rev, accompanied by an internal ribosomal entry site (IRES) for translation, upon HIV-1 infection (106). To obtain the EC_50_ values, dose-dependence data was plotted in GraphPad Prism and analyzed with the log(compound concentration) compared to normalized fluorescence *via* a response–variable slope equation and averaged. TZM-GFP cells were used for cytotoxicity, and cell viability was assessed with an XTT kit (Roche) according to the manufacturer’s instructions then CC_50_ values were determined using the same method as the EC_50_ values. TZM-GFP cells (HRP-20041) and the HIV-1_NL4-3_ virus (ARP-3418) were gifts from the NIH HIV Reagent Program.

### Endogenous Reverse Transcription (ERT) Assay

The ERT assay was used to determine the levels of reverse transcription products within native HIV-1 capsid cores, protocol adapted from (32, 49). Pseudotyped HIV-1 virions were produced in HEK-293/17 cells, as mentioned above. As previously reported (14), virions were permeabilized with melittin, then p24 levels for the sample was determined by ELISA using HIV-Ig (ARP-3957) as the primary antibody and anti-human HRP (Rockland Immunochemicals) as the secondary antibody, followed by quantification by a GloMax Navigator Plate Reader (Promega) after the chemiluminescent substrate (ThermoFisher) was added.

Purified virions, normalized to 500 ng p24, were treated with TURBO DNA-*free*™ DNase (AM1907), and various concentrations of metabolites were added in the presence of 10 mM Tris-HCl (pH 7.8), 75 mM NaCl, 2 mM MgCl_2_ 2.5 μg/mL melittin, 80 μM IP6, 0.5 mg/mL BSA, and 40 µM dNTPs at 37°C for 16 h. The reaction steps were quantified by qPCR with SYBR green detection using the following primer sets: 5′-GAGCTCTCTGGCTAACTAG-3′ and 5′-TGACTAAAAGGGTCTGAGGGATCT-3′ (minus-strand strong stop), 5′-GAGCCCTCAGATGCTGCATAT-3′ and 5′-CCACACTGACTAAAAGGGTCTGAG-3′ (first-strand transfer), 5′-TGTGTGCCCGTCTGTTGTGT-3′ and 5′-GAGTCCTGCGTCGAGAGAGC-3′ (second-strand transfer) adapted from (49).

### Reverse Transcriptase Primer Extension

Primer extension reactions were perform as previously described (70, 71). A Cy3-labled primer was annealed to the template at 1:2 molar ratio. Gallic acid and control inhibitors were incubated with RT for 30 minutes. Then, reactions were initiated by adding master mix containing 20 nM annealed Primer-Template substrate, 15 µM dNTPs, 6 mM MgCl2 in RT buffer (50 mM Tris, pH 7.8 and 50 mM NaCl) to a final volume of 20 µL and final RT concentration of 100 nM. The reactions were terminated after 60 minutes by adding an equal volume of 100% formamide containing bromophenol blue. Reaction products were resolved on a 7 M urea / 15% polyacrylamide gel, followed imaging on a phosphoimager (Typhoon FLA 9500).

The primer used are 5-Cy3-GTCACTGTTCGAGCACA-3’ (18mer); template sequence is 5’-CCATAGCTAGCATTGGTGCTCGAACAGTGAC-3’ (31mer), from (70).

### CA assembly assay and Transmission Electron Microscopy (TEM)

The *in vitro* CA assembly assay uses optical density as a proxy for lattice assembly (68), and these were conducted as previously reported (14, 68). Concentrated WT CA was diluted to 100 μM in 50 mM Tris (pH 8.0), treated with 100 μM of compound or vehicle control (DMSO or dH2O), and incubated on ice for at least 30 minutes. Then, to initiate assembly, high-salt buffer was added at a 1:1 ratio to the compound-treated sample in a 96-well plate. Each well contained 100 µL of 50 μM WT CA, 50 μM compound, and 2 M NaCl in 50 mM Tris (pH 8.0). Data were collected on a BioTek Neo plate reader, with A_350_ measured every 25 s for 2.5 h at room temperature. Simultaneously, a background control was measured that contained 50 μM compound and 50 μM WT CA in 50 mM Tris, with no NaCl; measurements of the protein assembly had the background readings subtracted. GraphPad Prism was used for visualization and statistical analysis.

The negative stain TEM protocol was adapted from (48). Briefly, 500 μM WT CA in 50 mM MES (pH 6.2) was treated with 500 µM metabolite (1:1), then capsid-like particles (CLPs) were assembled by mixing a 1:1 volume of the preincubated WT CA with a buffer that contains an excess of IP6 at 37°C. See (14, 48, 50).

## Acknowledgments

We would like to thank the members of the Sarafianos lab for feedback during TSAR package development. This project was supported in part by the National Institutes of Health (R01 AI120860 and U54 AI170855 to S.G.S.; F31 AI174951 to W.M.M.; F31 AI179424 to X.W.; W.M.M. and Z.C.L. were supported in part by T32 GM135060). The content is solely the responsibility of the authors and does not necessarily represent the official views of the National Institutes of Health. S.G.S. acknowledges funding from the Nahmias-Schinazi Distinguished Chair in Research. The plasmid encoding the Disulfide Stabilized CA_HEX_ (CA121) was kindly provided by Dr. Owen Pornillos (University of Utah); the plasmid encoding WT CA monomers was kindly provided by Dr. Robert Dick (Emory University); the plasmid encoding the 6HRT construct was kindly provided by Dr. Michael Parniak (University of Pittsburgh). LEN, TZM-gfp, HIV-Ig, and plasmid for pseudotyped virions were gifts from the NIH HIV Reagent Program. Electron Microscopy was carried out by Emory University Robert P. Apkarian Integrated Electron Microscopy Core Facility (RRID: SCR_023537). For providing essential details in making an R package, we would like to acknowledge the book R Packages by Hadley Wickham (O’Reilly) ©2015 Hadley Wickham, ISBN: 978-1-491-91059-7.

## Author contributions

Conceptualization, WMM, KAK, and SGS; Software, WMM, XG, and ZY; Methodology & Investigation, WMM, XG, ZY, XW, ZCL, HZ, JF and AE; Resources, WMM, ZCL, JF, AE and KAK; Visualization, WMM, XG, ZY, XW; Writing – Original Draft, WMM, XG, ZY and SGS; Writing – Review & Editing, WMM, XG, ZY, and SGS; Supervision, WMM, KAK, and SGS; Funding Acquisition, WMM, KAK and SGS.

---

**Supplementary Figure 1:**
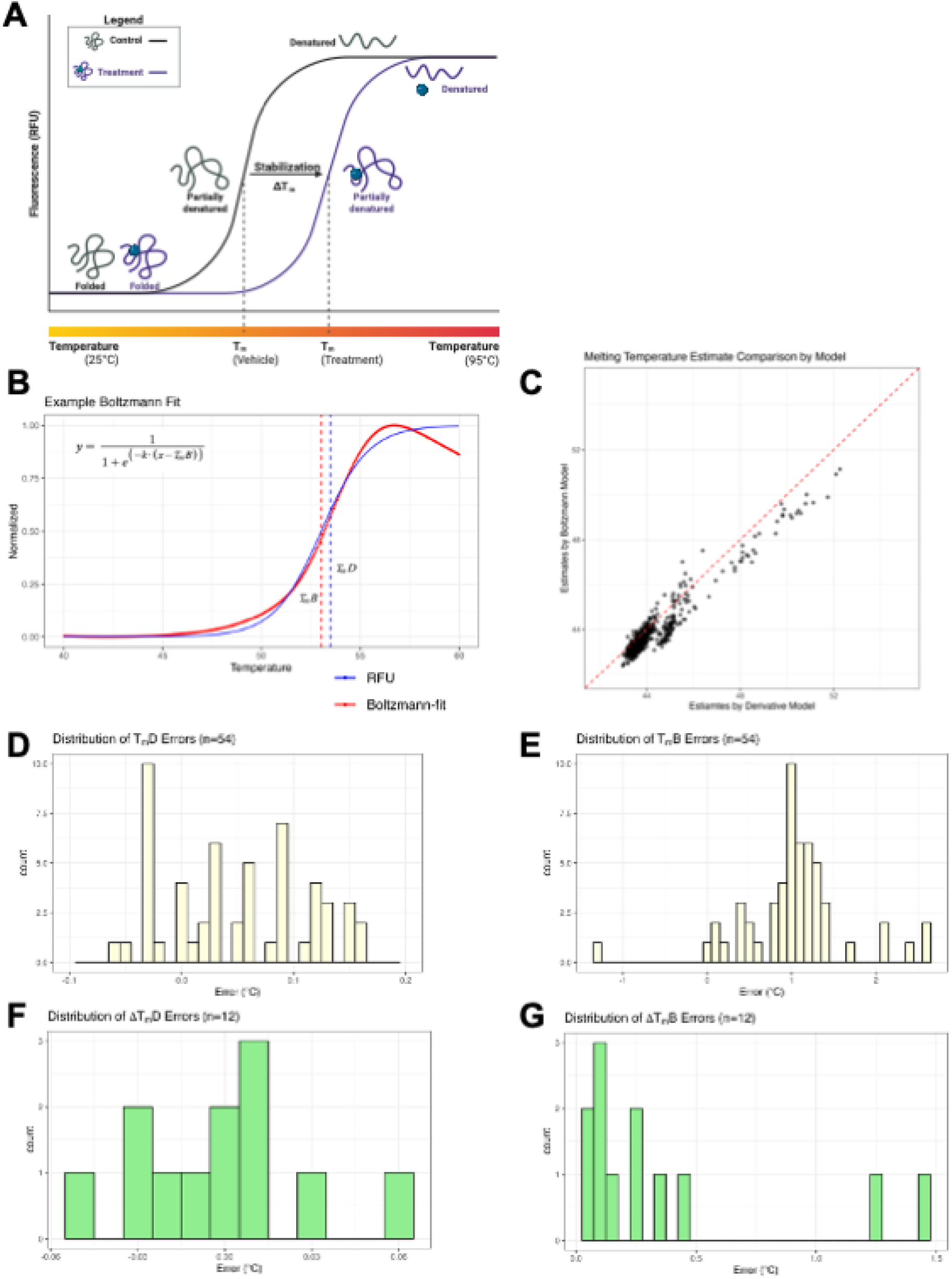
TSAR models and accuracy. **A)** Cartoon of thermal shift curves and protein denaturation to determine the 50% melting point (T_m_), change in T_m_ (ΔT_m_) from ligand-bound (blue circle) bound to the protein, thus increasing the temperature needed to denature the sample. Made with BioRender. **B)** Theoretical curves to exemplify the two modeling strategies between derivative T_m_ estimation (T_m_D), and Boltzmann T_m_ estimation (T_m_B). T_m_B uses the equation y=1/(1+*e*^−*k*(*x*−*x*2)^) (100). **C)** T_m_D vs T_m_B, showing overall correlation of the two values near the diagonal. **D, F)** Derivative estimation errors for the vitamin library. (n = 54) is N∼(0.048, 0.063). ΔT_m_D estimation error is similarly distributed, N∼(−0.001, 0.029). **E, G)** Boltzmann estimation errors (n = 54) is approximately N∼(1.029, 0.634). ΔT_m_B estimation error is biased in the positive direction, N∼(0.376, 0.451).

**Supplementary Figure 2:**
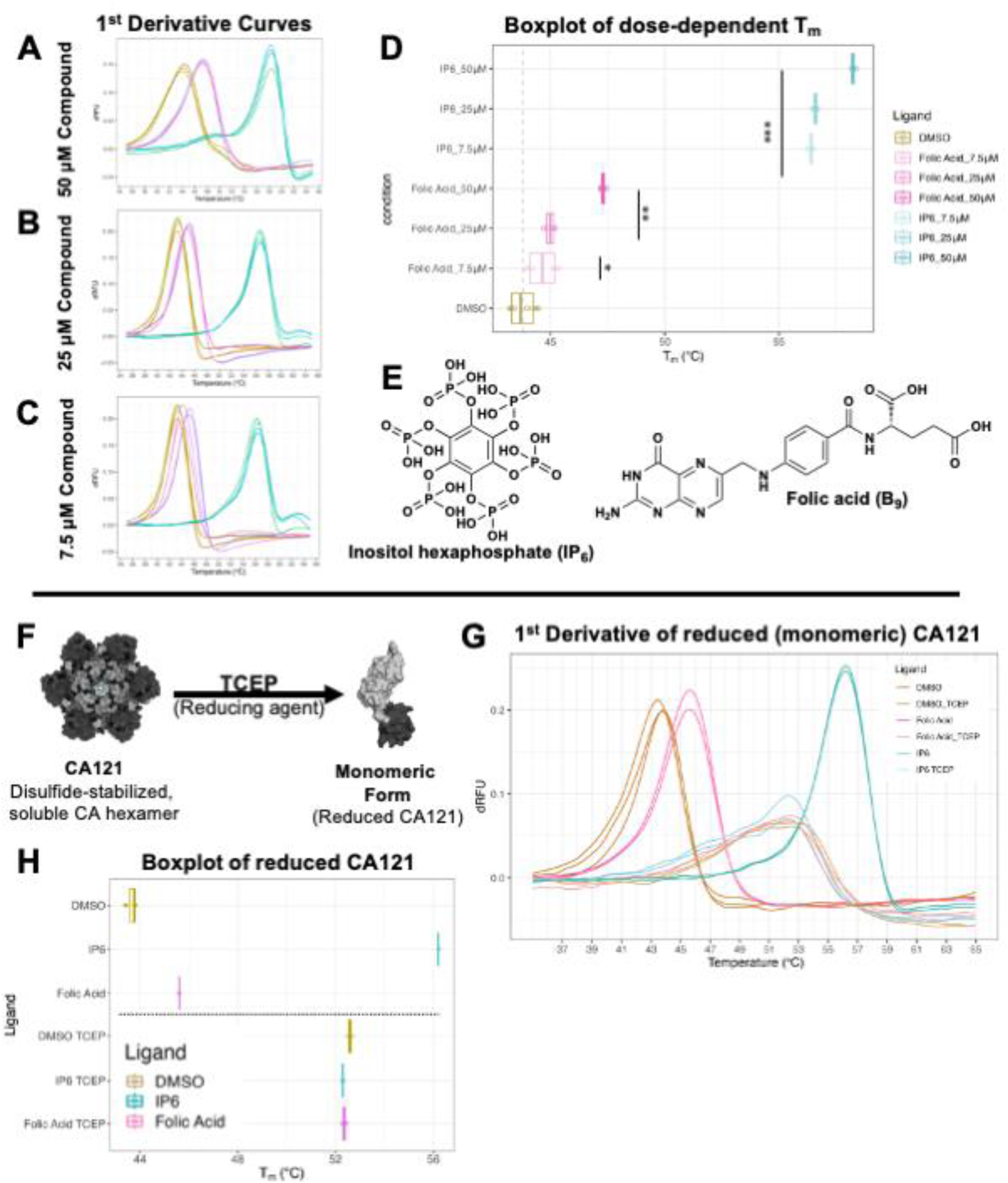
Dose dependence study of folic acid and CA_HEX_ shows ΔT_m_ is dosage-dependent. **A-C)** First derivative comparison graphs demonstrate that ΔT_m_ of folic acid with 7.5 µM CA121 (CA_HEX_) is dose dependent at **A)** 50 μM, **B)** 25 μM, **C)** 7.5 μM compound. D) TSAR-generated boxplot of (A-C), with one-tailed t-tests show that all treatment conditions are significant (*=p≤5E-3, **=p≤5E-5, and ***=p≤5E-9). **E)** Chemical Structures of hits, Folic Acid and IP6. Made in ChemDraw (V19.0). **F-H)** Reducing cross-linked CA121 shows thermal shift of folic acid requires hexameric status. **F)** TCEP was added as a reducing agent to break apart CA121, theoretically eliminating the central pore of protein-ligand interaction. **G)** TSA performed with ligands at 25 μM. Comparing first derivatives show that TCEP alters regular protein fluorescence curve. **H)** Boxplot demonstrates adding TCEP leads to insignificant ΔT_m_D.

**Supplementary Figure 3:**
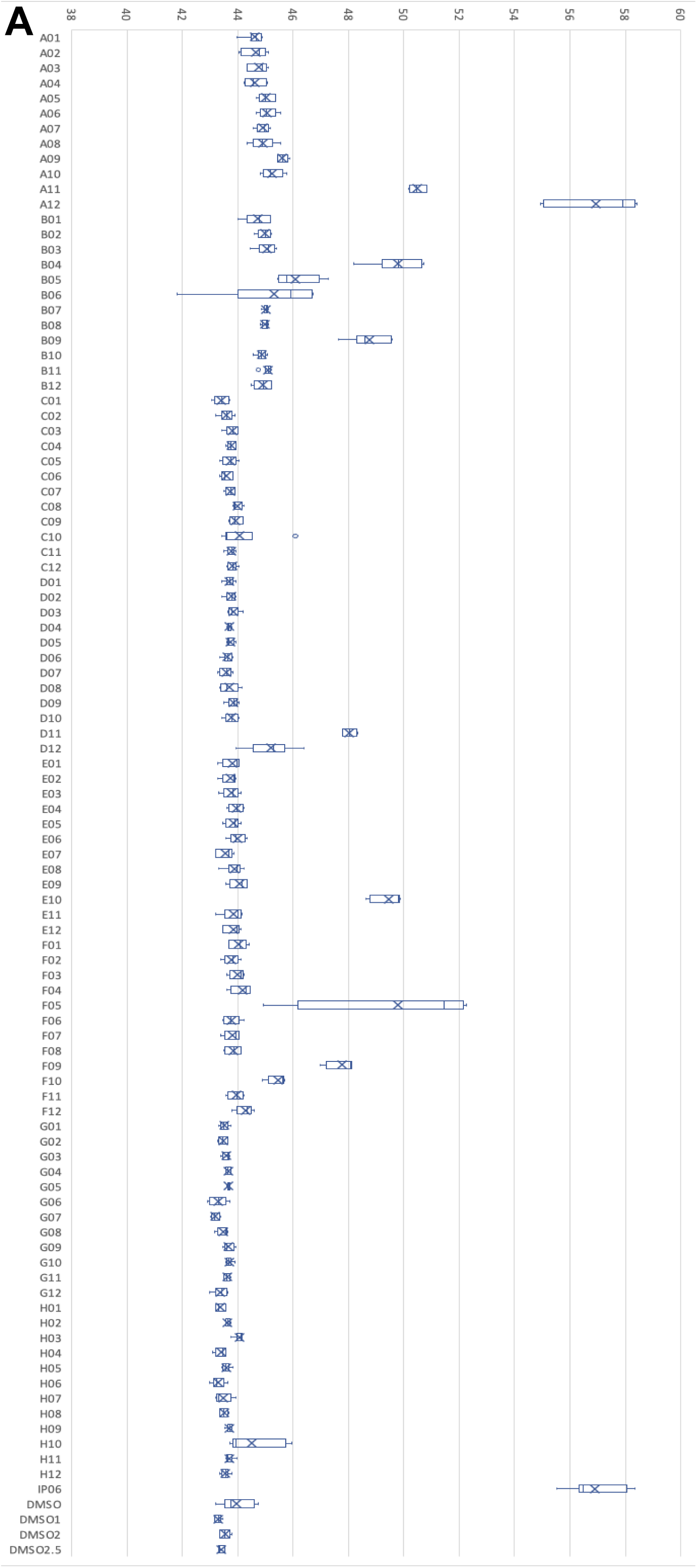
All hits from the 96-well organic metabolic acid library.

**Supplementary Figure 4:**
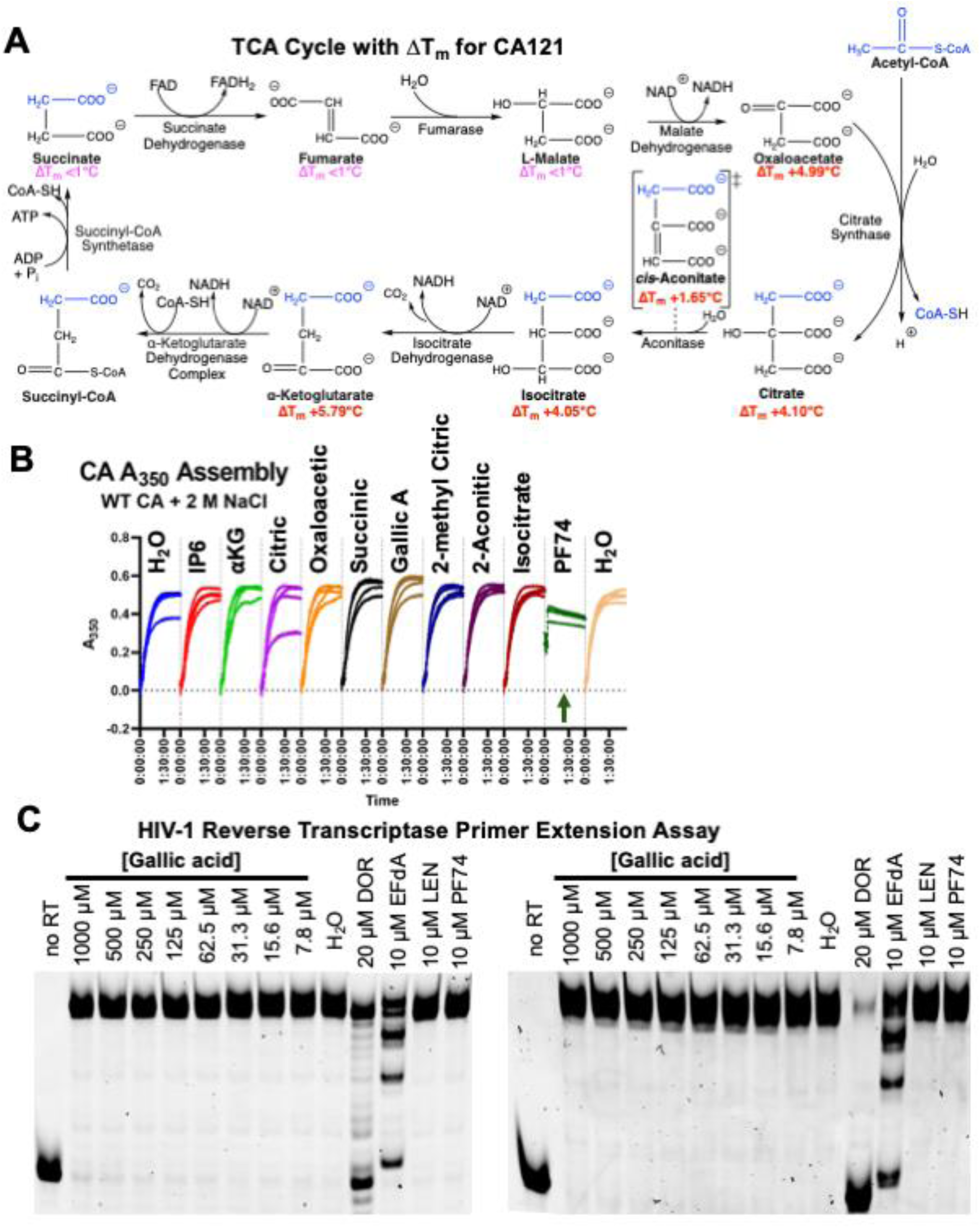
TCA cycle and *in vitro* HIV-1 assays testing metabolite hits. **A)** The tricarboxylic acid (TCA) cycle is shown, with T_m_ determined by TSAR labeled below for those tested against CA_HEX_. Made in ChemDraw (V19.0). **B)** An *in vitro* assembly assay for CA proteins that uses turbidity (A_350_) as a proxy for determining the rate of lattice formations (14, 68). The presence of metabolites does not seem to impact the rate of assembly. **C)** Testing the *in vitro* polymerase activity of HIV-1 RT using a primer-extension assay (70) finds that gallic acid does not inhibit the elongation of a DNA template. PF74 and LEN are CA-targeting inhibitors, and DOR and EFdA are RT-targeting inhibitors.

